# Lapidary: Identifying and reporting amino acid sequences in metagenomes using sequence reads and Diamond

**DOI:** 10.1101/2024.03.25.586564

**Authors:** Samuel J Bloomfield, Aldert L Zomer, Alison E Mather

**Author notes:** Correspondence,: Samuel J Bloomfield,; Alison E Mather.

## Abstract

Genome and metagenome comparisons rely on identifying genetic elements that differ or are in common between samples. These genetic elements can be identified by assembling sequenced reads and identifying the genetic element in the assembly, or by aligning nucleotide sequences in the reads to the nucleotide sequences of a reference genetic element. The first relies on the complete assembly of the genetic element of interest, and the second relies on a reference sequence represented in nucleotides. This is particularly challenging with metagenome data, where the genetic elements, including genes, are often fragmented because sequences are shared between different species in the metagenomic data, resulting in contig breaks in or around genetic elements. This presents a difficulty when identifying genetic elements through the first approach. A common approach with metagenomes is to map reads against reference nucleotide sequences and extract the depth and coverage from those reference sequences. However, currently no software exists to identity and report genetic elements using DNA-protein alignments in metagenomes. We have developed the software Lapidary to identify the identity, coverage, depth, and most likely sequence of amino acid sequences from both genome and metagenome read files. We tested the effectiveness of the method against simulated, genomic and metagenomic read datasets. Lapidary is more sensitive than assembly methods for metagenomic data that often have fragmented assemblies but is less sensitive when assemblies are more complete, as is the case with genomic data.

## Introduction

Genome and metagenome comparisons rely on identifying genetic elements that differ or are in common between samples [1, 2]. This requires software that identifies the genetic element in all samples with high sensitivity and specificity. Software programmes such as Prokka [3] and Bakta [4] can annotate nucleotide sequences to identify amino acid sequences [3] and programmes such as tBLASTn search for specific amino acid sequences [5]. However, aligning nucleotide sequences to amino acid sequences can be computationally intensive as nucleotide sequences are read as three-base codons during translation and can be read in the forward and reverse directions, leaving six open reading frames that need to be considered. It is also difficult to determine which open reading frames encode for an amino acid sequence, particularly for short amino acid sequences [6]. These programmes also rely on completely assembled elements and struggle with metagenomic assemblies where due to the number of similar microorganisms, assemblies break at the repetitive regions, leading to fragmented assemblies consisting of many short contigs.

DNA sequences of interest can also be identified by aligning sequenced reads to reference genetic sequences using various alignment programmes [7–9]. Programmes have been developed to search a large number of nucleotide sequences for amino acid sequences [10]. However, no such programme has been developed to rapidly identify and investigate the depth and coverage of amino acid sequences of interest by read alignment. DNA-protein alignments are more sensitive because of higher levels of conservation in protein sequences [11], so such a programme could identify more related sequences than programmes that rely on a nucleotide database. In this manuscript we describe the software Lapidary, that aligns short reads of nucleotides to amino acid sequences to predict their coverage, identity, sequence depth and most likely sequence and apply the software to genomic, metagenomic and simulated datasets to test its sensitivity and specificity. Lapidary is the process of transforming gemstones into decorative items and was used to name the software as it relies on Diamond [12] for read alignments.

## Methods

### Lapidary

Lapidary uses Diamond v2.1.6 [12] to align short reads to an amino acid database using the BLOSUM45 scoring matrix [13] and extract the translated section of the read alignment (Figure 1). Reads are discarded if neither of their ends align to the amino acid sequence. For each amino acid sequence, the number of amino acids to which reads align is used to calculate the amino acid coverage and read depth. At each position, the best match is called from the read with the highest identity value according to Diamond to give the most likely sequence at that position. The number of matches between the most likely sequence and the reference is used to calculate the genetic identity. If reads align to multiple, separate sections of an amino acid sequence, calculations for coverage, depth and identity are performed for the longest amino acid section to which reads align.

**Figure 1.**
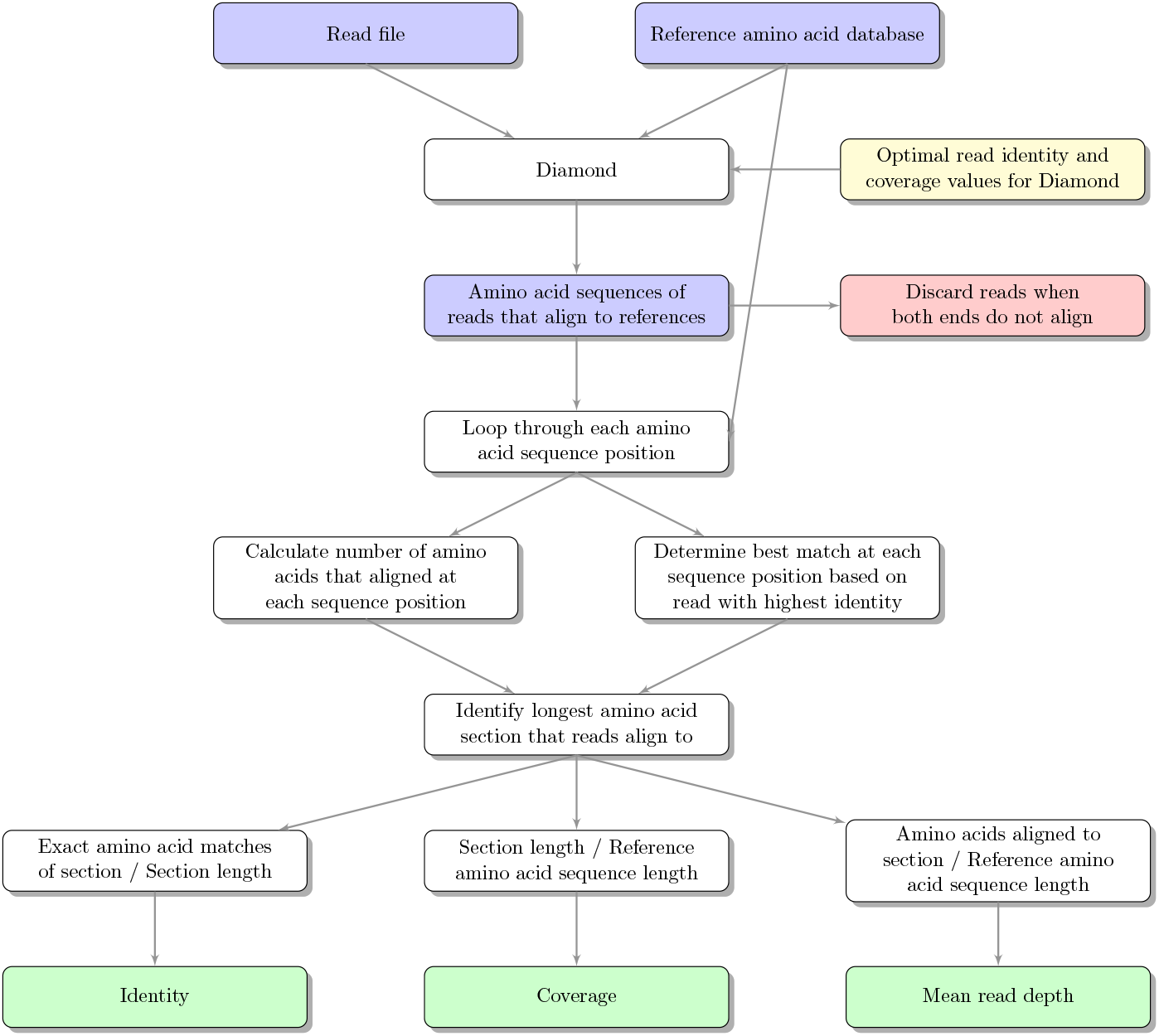
Lapidary workflow.

### Diamond parameter optimisation

BacMet is a database of amino acid sequences responsible for antibacterial biocide and metal-tolerance resistance [11]. Most of the amino acid sequences originate from bacteria that are easy to culture. To optimise the Diamond parameters for Lapidary, multiple simulated datasets of the amino acid sequences in the BacMet database were created by both reducing the length of the amino acid sequences whilst keeping the first and last amino acid (to alter the coverage calculations) and by randomly exchanging amino acids for others (to alter identity calculations). The coverage and identity values of the amino acid sequences in the permuted BacMet datasets varied from 50-100% length of the original sequences in increments of 10%, generating 36 datasets with known coverages and identities compared to the original dataset. Each of the simulated amino acid sequences in each of the 36 datasets were then converted to the most likely nucleotide sequence using the EMBOSS software v6.5.7 [14] backtranseq. ART v2.3.7 [15] was used to simulate 150 single-end reads from each of the amino acid datasets at a 10-fold read depth. The simulated reads were analysed with Lapidary with Diamond coverage read cut-offs ranging from 10-90% in increments of 10% and Diamond identity read cut-offs ranging from 50-90% in increments of 10% to identify the optimal combination of these parameters to maximise the sensitivity and specificity of the method. The cut-offs commonly used to identify genetic elements of interest (e.g., antimicrobial resistance genes) in metagenomes were used: 60% coverage and 90% identity [1]. If the simulated read dataset was formed from an amino acid dataset with 90% identity and 60% coverage or above to the original and Lapidary detected it, this was counted as a true positive, but if Lapidary failed to detect it this was considered a false negative. If the read dataset was formed from a simulated amino acid database with less than 90% identity or 60% coverage and Lapidary detected it this was counted as a false positive, but if Lapidary failed to detect it this was considered a true negative.

To test the effect of read depth, reads were simulated with a 100-fold read depth and Lapidary was used to detect amino acids sequences in the BacMet databases as above with the optimal Diamond parameters.

### Genome dataset

The raw Illumina reads of 68 previously published *Salmonella* genomes [2] were trimmed using Trimmomatic using 2:30:10 as the ILLUMINACLIP, 3 as the LEADING, 3 as the TRAILING, 4:25 as the SLIDINGWINDOW and 50 as the minimum length parameters. Paired trimmed reads were placed through Lapidary with the BacMet database. The trimmed reads were also assembled using SPAdes [16] in “careful” mode. These isolates had also been assembled with PacBio long reads using various hybrid assembly methods to obtain less fragmented assemblies. The hybrid assemblies of this previous study were obtained and along with the SPAdes assemblies, were searched for amino acid sequences belonging to the BacMet database using tBLASTn. Amino acid sequences with a 90% coverage and 90% identity were considered present for genomes [9]. For these genome datasets, an additional mean read depth of 10 was included for Lapidary.

### Metagenome dataset

The raw Illumina reads of seven previously published retail chicken meat metagenomes [1] were trimmed using Trimmomatic as above and placed through Lapidary with the BacMet database. The trimmed reads were also assembled using MegaHit [17]. The seven metagenomes had also previously been sequenced using the Oxford PromethION platform at a depth of 8Gb. The raw long-reads were trimmed using Nanofilt v0.1.0 (https://github.com/wdecoster/nanofilt). Trimmed long- and short-reads were assembled using OPERA-MS v0.9.0 [18]. tBLASTn was used to search the metagenome assemblies against the BacMet database. Amino acid sequences with a 60% coverage and 90% identity were considered present for metagenomes. The BacMet database is made up of metal-tolerance genes from bacteria that are easy to culture [11], so the coverage cut-off was reduced for metagenomes to identify metal-tolerance genes in any distantly related bacteria present.

## Results

### Diamond parameter optimisation

The optimal Diamond read parameters for Lapidary was a Diamond read identity of 70% and a Diamond read coverage of 50% (Figure 2), which had a mean sensitivity of 50.7% (0.8-96.1%) and a mean specificity of 99.9% (99.6-100%) with 10-fold read depth simulated reads. Increasing the read depth increased the mean sensitivity to 74.2% (16.6-96.8%) but had little effect on the mean specificity of 99.9% (98.9-100%) (Figure 3). Most Lapidary coverage and identity estimates were close to the actual values of the simulated datasets, but Lapidary often underestimated the coverage (Figure 4).

**Figure 2.**
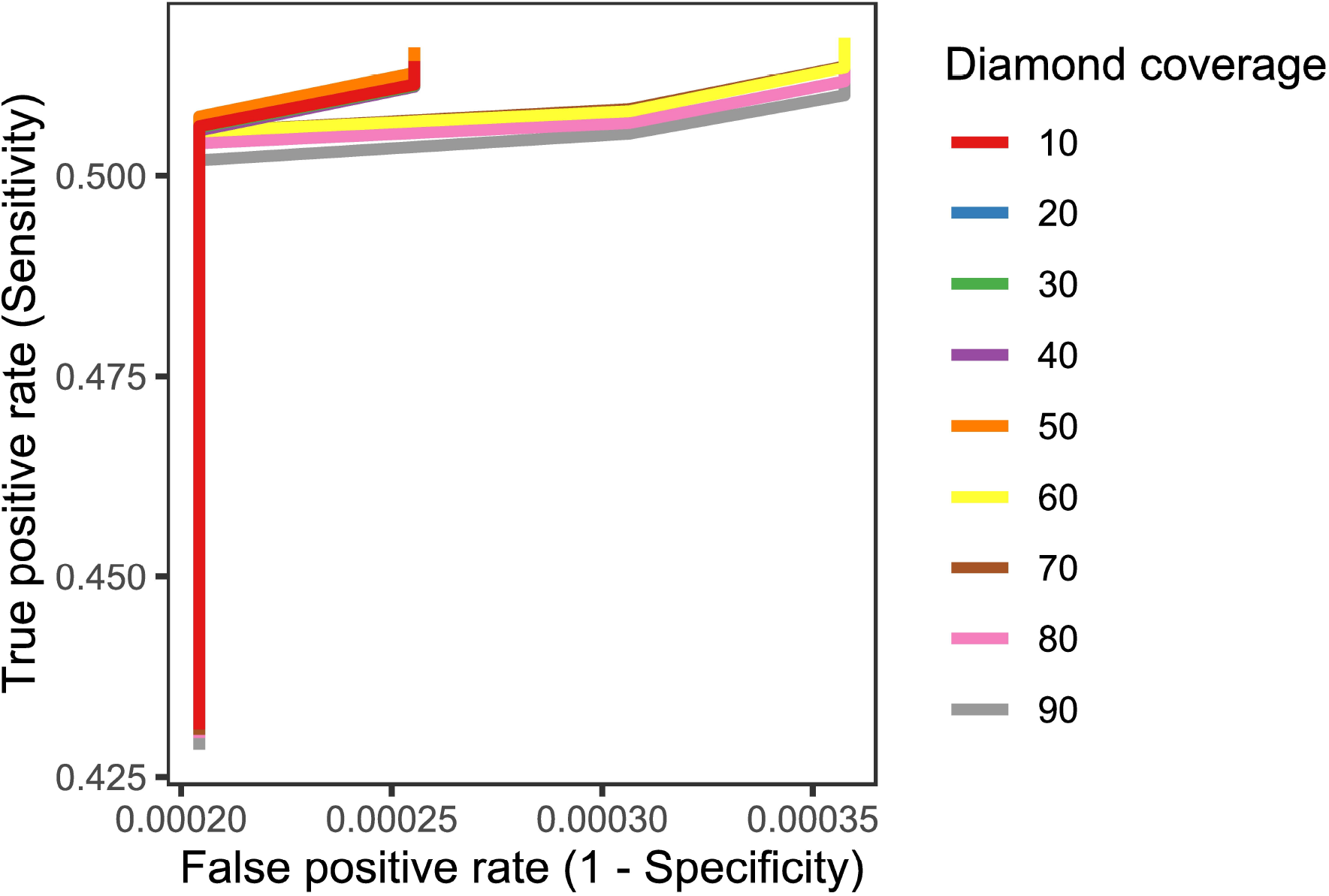
Sensitivity and specificity of Lapidary using different Diamond identity and coverage values on simulated datasets.

**Figure 3.**
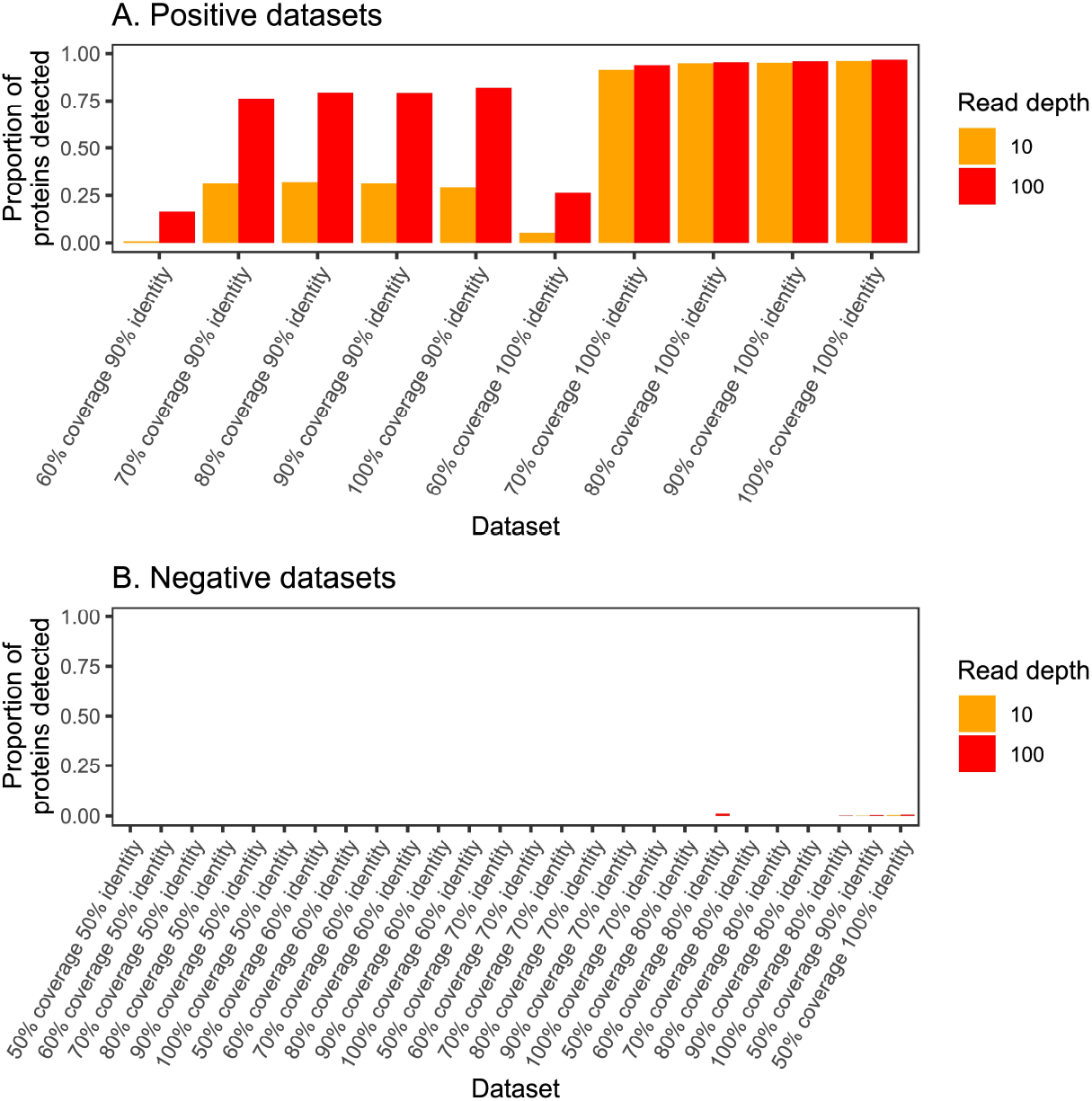
Proportion of amino acid sequences detected using different simulated read datasets that meet (A; positive datasets) and do not meet (B; negative datasets) the cut-offs and coloured by the read depth of these read datasets.

**Figure 4.**
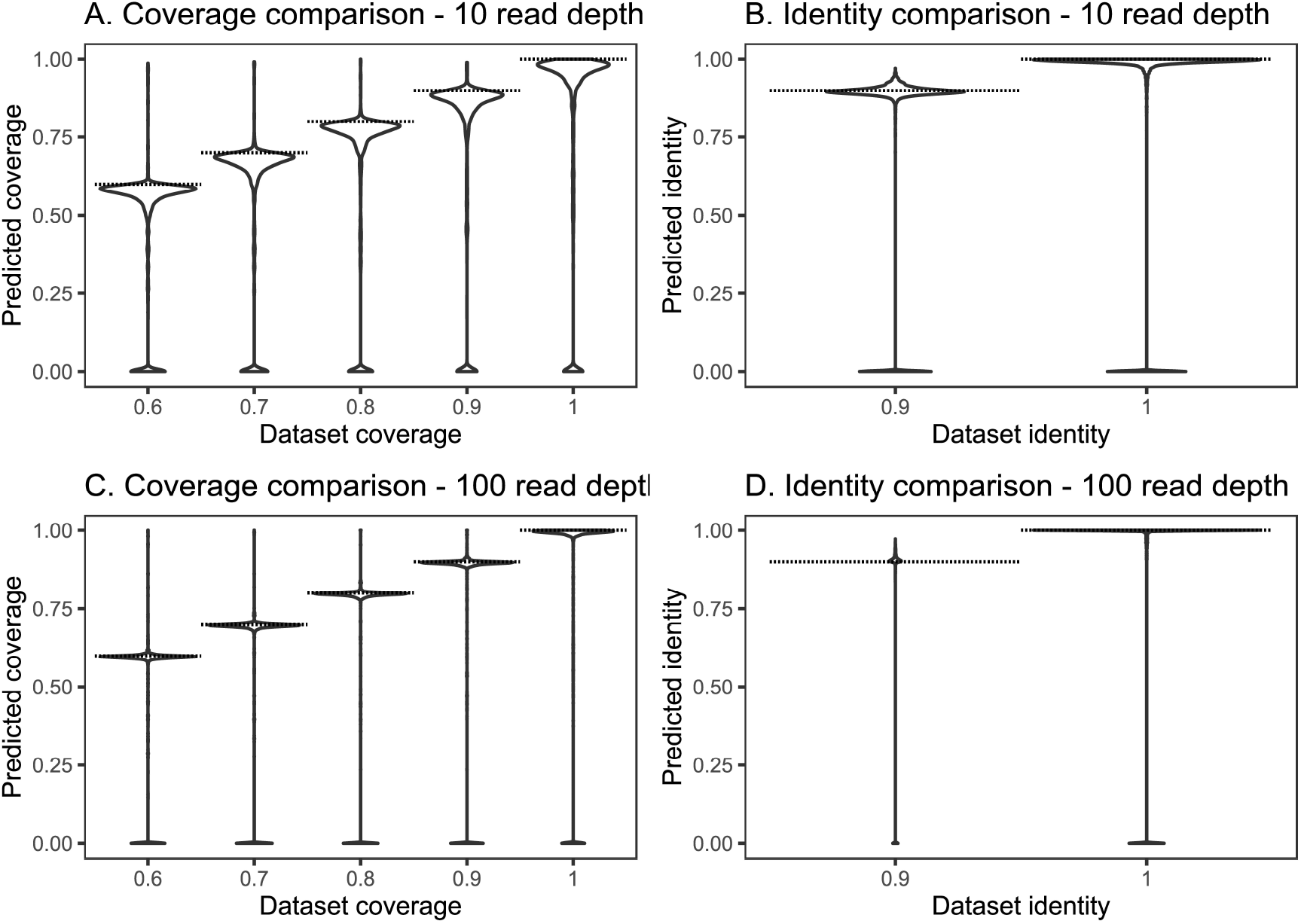
Lapidary coverage (A and C) and identity (B and D) estimates versus the actual values for simulated datasets (dashed lines) at 10-fold (A and B) and 100-fold (C and D) read depth.

### Genome dataset

Using a combination of Lapidary and tBLASTn on the short-read (Short) and hybrid (Long) assemblies of 68 *Salmonella* genomes, 85-132 metal-tolerance amino acid sequences were identified (Figure 5). Assuming that tBLASTn on short and long assemblies accurately identified all amino acid sequences (Table 1), Lapidary had a mean sensitivity of 92.3% (85.3-97.6%) and specificity of 95.2% (93.1-96.7%) in this genome dataset.

**Table 1.**
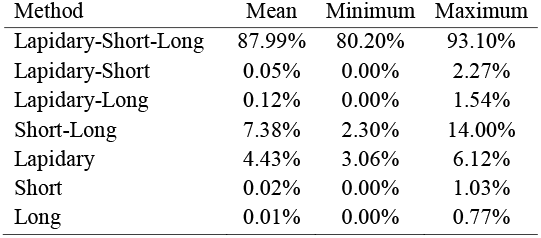
Percentage of metal-tolerance amino acid sequences identified in 68 *Salmonella* genomes using only Lapidary, or tBLASTn using short-read (Short) or hybrid (Long) assemblies, or by multiple methods (Lapidary-Short, Lapidary-Long, Short-Long, Lapidary-Short-Long).

**Figure 5.**
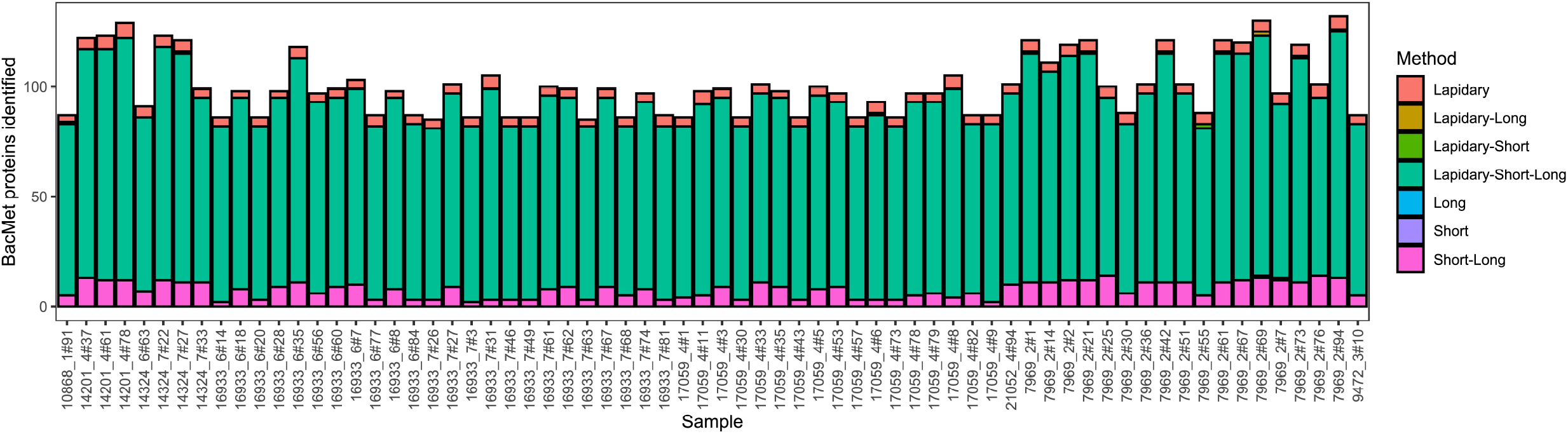
Number of metal-tolerance genes identified in 68 *Salmonella* genomes using only Lapidary or tBLASTn on short-read (Short) or hybrid (Long) assemblies, or by multiple methods (Lapidary-Short, Lapidary-Long, Short-Long, Lapidary-Short-Long).

### Metagenome dataset

Using a combination of Lapidary and tBLASTn on the short-read (Short) and hybrid (Long) assemblies of seven retail chicken metagenomes, 44-153 metal-tolerance amino acid sequences were identified (Figure 6). Overall, Lapidary identified 21.4-70.9% more metal-tolerance genes compared to tBLASTn (Table 2).

**Table 2.** Percentage of metal-tolerance amino acid sequences identified in seven retail chicken metagenomes using only Lapidary, or tBLASTn using hybrid (Short) or long-read (Long) assemblies, or by multiple methods (Lapidary-Short, Lapidary-Long, Short-Long, Lapidary-Short-Long).

| Method | Mean | Minimum | Maximum |
| --- | --- | --- | --- |
| Lapidary-Short-Long | 57.01% | 46.43% | 66.67% |
| Lapidary-Long | 2.44% | 0.00% | 4.20% |
| Lapidary-Short | 6.11% | 2.52% | 8.33% |
| Short-Long | 2.84% | 0.00% | 4.76% |
| Lapidary | 29.93% | 17.65% | 41.50% |
| Long | 1.45% | 0.00% | 4.20% |
| Short | 0.22% | 0.00% | 0.86% |

**Figure 6.**
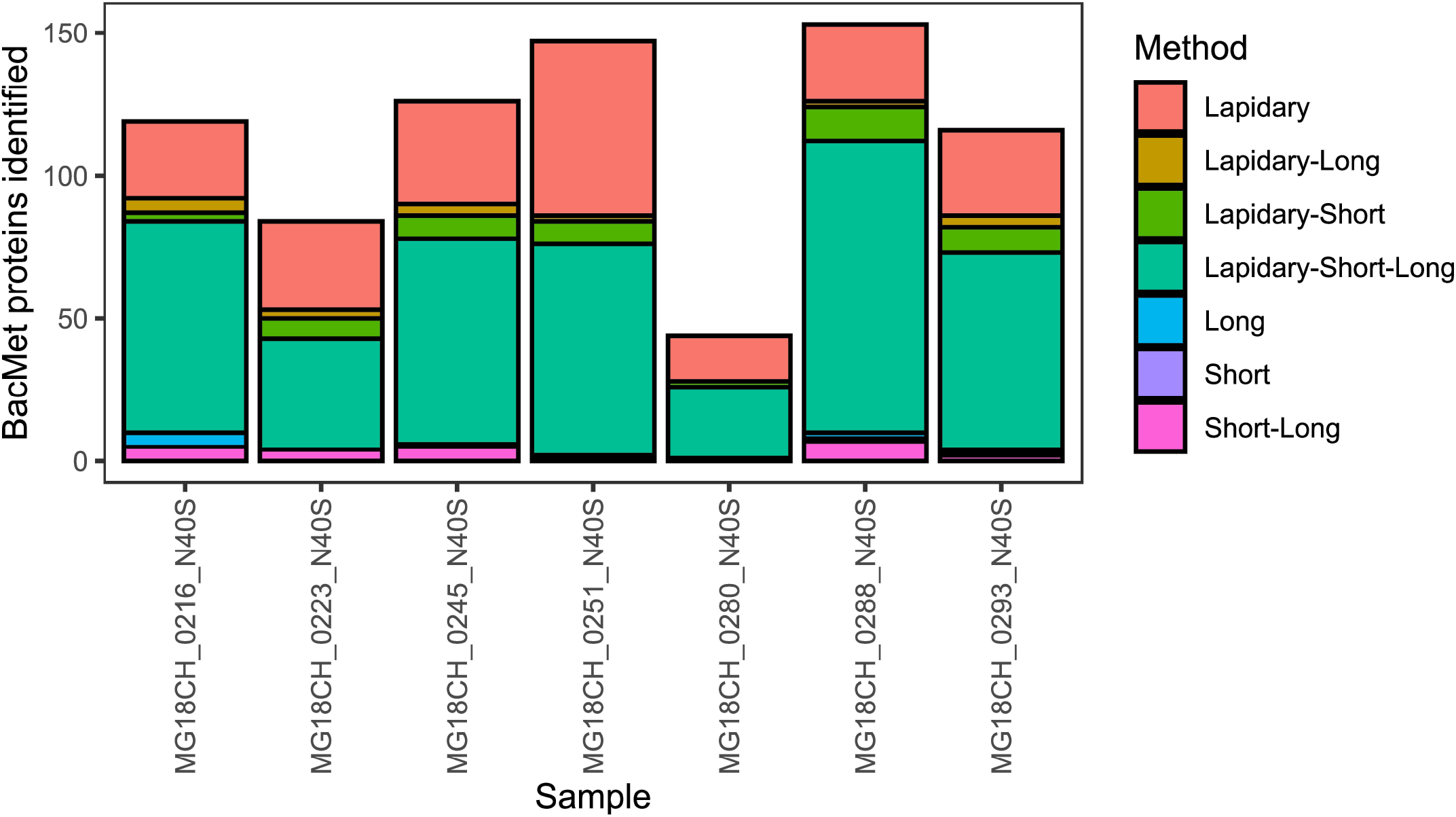
Number of metal-tolerance genes identified in seven retail chicken metagenomes using only Lapidary or tBLASTn on short-read (short) or hybrid (long) assemblies, or by multiple methods (Lapidary-Short, Lapidary-Long, Short-Long, Lapidary-Short-Long).

## Discussion

Lapidary allows the identification of amino acid sequences using short-read sequences, and it reports the identity, coverage, depth, and most likely amino acid sequence, but its effectiveness varied with simulated, genomic and metagenomic reads. With simulated reads the method was very specific but was not very sensitive when the original dataset used to simulate the reads had a coverage identical to the cut-off used for identifying amino acid sequences. This can be attributed to the mismatches artificially added to the BacMet database to create the simulated databases. These mismatches were not uniformly distributed, resulting in some amino acid sequences having 90% identity over the entire sequence, but sections of the amino acid sequence with much lower identity. Lapidary will not be able to align reads to these low identity regions, even with the BLOSUM45 scoring matrix to allow for the identification of distant homologs [13], preventing the software from identifying the amino acid sequences; this explains why Lapidary often underestimated the coverage. Decreasing the Diamond read identity cut-off could help account for this but will also result in reads from unrelated amino acid sequences matching, which will decrease specificity.

Lapidary sensitivity was higher for the genome dataset (92.3%) compared to the simulated dataset (74.2%), but the specificity was lower for the genome dataset (95.2%) compared to the simulated dataset (99.9%). The higher sensitivity of the genome dataset is likely due to this dataset containing less protein sequences on the coverage and identity cut-offs, whilst the lower specificity can be attributed to genomes containing reads that encode for a wide range of amino acid sequences, some of which may be similar to sections of the reference amino acid sequences, resulting in their false identification. Lapidary could be modified to increase the specificity for the genome data by local assembly of the aligned reads similar to the process used in ARIBA [19], but this would not be applicable for metagenome data where the presence of reads from multiple microorganisms would limit the assembly step.

Application of Lapidary to metagenome reads identified more amino acid sequences than were detected by assembling the reads and identifying the amino acid sequences using tBLASTn. This is because metagenome assemblies contain fragmented regions, even those assembled with long-reads, and tBLASTn relies on the amino acid sequence to be complete in order to identify it. However, better assemblers are being developed to create less fragmented metagenome assemblies [20]. In the meantime, Lapidary can identify more amino acid sequences than traditional assembly and tBLASTn methods for metagenomes.

Metagenome datasets may contain reads that originate from multiple similar genes compared to a reference, but here we are interested in the most likely sequence even if it is not the most prominent. For this reason, Lapidary calls the most likely sequence based on the reads with the highest identity to the reference sequence as calculated by Diamond. For genome datasets there may be a small amount of contamination in which we are not interested, but this may obscure results. A consensus sequence would prevent small amounts of contamination influencing results. Although we based all calculations in this study on reads with the highest identity, Lapidary has the option to calculate and use a consensus sequence for its calculations.

Lapidary was optimised for 150 bp paired-end Illumina reads. Application of Lapidary to different read sets will need the Diamond coverage and identity values to be optimised for different sized reads. This study also did not take into consideration the quality of the reads; this might affect read length, if low quality reads are trimmed, and may also influence the optimal Diamond parameters. However, we have described here a method to optimise Lapidary for whatever read dataset is at hand.

## Conclusion

Lapidary is a software that can identify amino acid sequences from raw nucleotide reads and estimate the most likely amino acid sequence, either from genomes or metagenomes. This type of tool was not previously available, given the difficulty of aligning nucleotide reads to amino acid sequences. It is less sensitive than tBLASTn when assemblies are more complete, as are generally obtained with whole genome sequences of bacteria, but more sensitive than tBLASTn for metagenome data where assemblies are often fragmented.

## Abbreviations

BBSRC: Biotechnology and Biological Sciences Research Council;
BLAST: Basic Local Alignment Search Tool;
ENA: European Nucleotide Archives;
FSA: Food Standards Agency;
SRA: Sequence Read Archive.

## Impact statement

Genome and metagenome comparisons rely on identifying genetic elements that differ or are in common between samples. Identifying amino acid sequences usually relies on assembling sequence reads and then searching the assembly for the amino acid sequence. Metagenomes often contain very fragmented assemblies, limiting the identification of many amino acid sequences. We developed the software Lapidary to identify the identity, coverage, depth, and most likely sequence of amino acid sequences without the need for read assembly. Lapidary is more sensitive than assembly-based methods for metagenomic data that often have fragmented assemblies but is less sensitive when assemblies are more complete, as is the case with genomic data.

## Data summary

Genome short and long reads were previously uploaded to the European Nucleotide Archive (ENA) under projects PRJEB1397, PRJEB9121 and PRJEB2973 for Illumina, and PRJEB9562 for PacBio data.

Metagenome short reads were previously uploaded to the Sequence Read Archive (SRA) under project PRJNA849983. Metagenome long-reads were uploaded to SRA under project PRJNA1034280.

The Lapidary software is open source and available for Linus at GitHub under the GNU GPLv3 license: https://github.com/samuelbloomfield/lapidary.

A platform-independent web interface for Lapidary is available: https://lapidary.quadram.ac.uk/.

## Author statements

### Author contributions

Conceptualisation, SJB, ALZ, AEM; Methodology, SJB and ALZ; Writing-original draft preparation, SJB; Writing -review and editing, SJB, ALZ and AEM; Supervision, AEM.

## References

1. Bloomfield SJ, Zomer AL, O’Grady J, Kay GL, Wain J, et al. Determination and quantification of microbial communities and antimicrobial resistance on food through host DNA-depleted metagenomics. Food Microbiol 2023;110:1–12.

2. Bloomfield S, Duong VT, Tuyen HT, Campbell JI, Thomson NR, et al. Mobility of antimicrobial resistance across serovars and disease presentations in non-typhoidal Salmonella from animals and humans in Vietnam. Microb Genomics 2022;8:1–13.

3. Seemann T. Prokka: rapid prokaryotic genome annotation. Bioinformatics 2014;30:2068–2069.

4. Schwengers O, Jelonek L, Dieckmann MA, Beyvers S, Blom J, et al. Bakta: rapid and standardized annotation of bacterial genomes via alignment-free sequence identification. Microb Genomics 2021;7:1–13.

5. Gertz EM, Yu Y-K, Agarwala R, Schaffer AA, Altschul SF. Composition-based statistics and translated nucleotide searches: Improving the tBLASTn module of BLAST. BMC Biol 2006;4:1–14.

6. Ansong C, Purvine SO, Adkins JN, Lipton MS, Smith RD. Proteogenomics: needs and roles to be filled by proteomics in genome annotation. Brief Funct Genomics 2008;7:50–62.

7. Li H, Durbin R. Fast and accurate short read alignment with Burrows-Wheeler transform. Bioinformatics 2009;25:1754–1760.

8. Inouye M, Dashnow H, Raven L, Schultz MB, Pope BJ, et al. SRST2: Rapid genomic surveillance for public health and hospital microbiology labs. Genome Med 2014;6:1–16.

9. Clausen PTLC, Aarestrup FM, Lund O. Rapid and precise alignment of raw reads against redundant databases with KMA. BMC Bioinformatics 2018;19:1–8.

10. de Weerd H, van der Veen BE, Claesson MJ. BlastXtract2: Improving early exploration of (meta) genomes. Bioinformation 2015;11:173–175.

11. Pal C, Bengtsson-Palme J, Rensing C, Kristiansson E, Larsson DGJ. BacMet: antibacterial biocide and metal resistance genes database. Nucleic Acid Res 2014;42:D737–D743.

12. Buchfink B, Reuter K, Drost H-G. Sensitive protein alignments at tree-of-life scale using DIAMOND. Nat Methods 2021;18:366–368.

13. Henikoff S, Henikoff JG. Amino acid substitution matrices from protein blocks. Proc Natl Acad Sci 1992;89:10915–10919.

14. Rice P, Longden I, Bleasby A. EMBOSS: The European Molecular Biology Open Software Suite. Trends Genet 2000;16:276–277.

15. Huang W, Li L, Myers JR, Marth GT. ART: a next-generation sequencing read simulator. Bioinformatics 2012;28:593–594.

16. Bankevich A, Nurk S, Antipov D, Gurevich AA, Dvorkin M, et al. SPAdes: a new genome assembly algorithm and its applications to single-cell sequencing. J Compulational Biol 2012;19:455–477.

17. Li D, Liu C-M, Luo R, Sadakane K, Lam T-W. MEGAHIT: an ultra-fast single-node solution for large and complex metagenomics assembly via succinct de Bruijn graph. Bioinformatics 2015;31:1674–1676.

18. Gao S, Bertrand D, Chia BKH, Nagarajan N. OPERA-LG: efficient and exact scaffolding of large, repeat-rich eukaryotic genomes with performance guarantees. Genome Biol 2016;17:102.

19. Hunt M, Mather AE, Sanchez-Buso L, Page AJ, Parkhill J, et al. ARIBA: rapid antimicrobial resistance genotyping directly from sequencing reads. Microb Genomics 2017;3:1–11.

20. Steinegger M, Mirdita M, Söding J. Protein-level assembly increases protein sequence recovery from metagenomic samples manyfold. Nat Methods 2019;16:603– 606.

